# Social, cognitive and sensory dimensions of cortical network overconnectivity in young children with autism spectrum disorder

**DOI:** 10.1101/2025.02.12.637813

**Authors:** Borja Rodríguez-Herreros, Ahmad Mheich, Joana Maria Almeida Osório, Sonia Richetin, Vincent Junod, Lorène Arnold, Victoria Aeschbach, Lilia Nora Adamou, Katherina Gschwend, Laura Mendes, David Romascano, Paola Yu, Marine Jequier Gygax, Anne M. Maillard, Micah Murray, Mahmoud Hassan, Nadia Chabane

## Abstract

The heterogeneity in both the neurobiological mechanisms and the phenotypic presentation of autism spectrum disorder (ASD) poses a major challenge to clinical and translational research. Early inefficiencies in functional connectivity (FC) have been associated with ASD, yet it remains unclear whether and how abnormal brain network properties may account for individual differences across ASD-related symptomatology and behaviors. We applied source-level reconstruction to resting-state high-definition EEG data in a cohort of 113 young children (40 with ASD) to identify early global and local alterations of cortical network connectivity. We subsequently used regularized canonical correlation analysis (rCCA) to characterize specific FC patterns linked to variation in cognitive, social and sensory dimensions within the autism spectrum. We found increased low-frequency FC in frontotemporal cross-hemispheric networks and lateral-occipital regions of young ASD children. RCCA revealed three distinct FC patterns in recurrent ASD-related networks, each contributing to predict individual differences in cognitive, social and sensory features. These linked FC-behavior dimensions may shed light on atypical brain network topology conferring risk for specific phenotypic manifestations of ASD, which may implicate unique underlying neurobiological mechanisms.

## Introduction

Autism spectrum disorder (ASD) is a neurodevelopmental disorder (NDD) classically characterized by persistent and significant social communication deficits and repetitive sensorimotor behaviors[1]. Understanding the early development of the neuronal circuitry underlying ASD is fundamental to elucidate its etiology and to promote more effective treatment interventions. The introduction of neuroimaging techniques has provided powerful means to investigate the neurophysiological underpinnings of ASD and other NDDs[2]. Over recent years, the identification of pathological functional and structural brain networks has become a topic of increasing interest, and it is considered one of the most promising prospects in the study of brain disorders[3]. Growing consensus suggests that disruptions in brain functional connectivity (FC) play an important role in the pathophysiology of ASD[4–6]. Several studies have reported that individuals with ASD exhibit abnormal FC, resulting in alterations of large-scale brain networks[7–11]. Yet, maturational changes of functional brain topology through early childhood and their association with clinically relevant ASD-related phenotypic presentations remain poorly understood.

From the clinical perspective of an as early as possible identification of brain functional alterations associated with ASD, the demand is high for non-invasive and easy-to-use methodological approaches. High-density resting-state electroencephalography (HD-EEG) has proven to be a practical and useful technique to explore large-scale brain physiological activity at rest in pediatric clinical populations[12, 13]. Specifically, the convenience of resting-state HD-EEG to monitor brain activity in the absence of overt task performance or sensory stimulation stands out compared to other functional imaging methods that are rather difficult to implement with young ASD children. A considerable number of studies have investigated atypical brain FC in ASD individuals using resting-state EEG protocols[14]. Of the wide repertoire of features that can be obtained, previous work has predominantly focused on the analysis of spectral power. These studies have reported inconsistent findings regarding altered brain connectivity in low (theta, alpha) and high frequency (gamma) band oscillations in ASD individuals[15]. Reduced theta power has been associated with greater ASD symptomatology[16], while local changes in alpha and gamma power have also been observed[17, 18]. Nevertheless, a relatively consistent profile of spectral power abnormalities in ASD children through development has emerged, suggesting excessive power in low and high frequency bands at early ages, but reduced power in the mid-range band[4, 19].

Besides spectral features, FC analysis has also been used to shed light on the brain network organization underpinning ASD. Several resting-state EEG studies revealed characteristic altered network topology in children diagnosed with both syndromic and non-syndromic ASD[20–23]. These studies support the developmental disconnection theory, which proposes a decreased long-range integration accompanied by local overconnectivity and reduced functional specialization in the brain[24]. Nevertheless, these EEG connectivity studies have all been performed in the sensor space (scalp level), thus using inflated connectivity estimates caused by volume conduction artefacts and preventing inferences about interacting brain regions[25, 26]. Only one study has conducted a source-level assessment to identify specific functional network properties associated with ASD[27], albeit with a markedly low-density EEG electrode placement. Therefore, disruptions of functional network topology in children with ASD have barely been explored with source-level HD-EEG. Moreover, whether altered network properties are reliably associated with the presence of specific clinical or ASD-related phenotypic manifestations is unclear. Efforts to associate clinical features with atypical brain network organization in ASD have been essentially addressed using resting-state functional MRI (fMRI), with controversial results likely due to ascertainment biases and the inherent etiological heterogeneity of ASD. For example, ASD core symptomatology and global cognition failed to be associated with several altered network metrics in ASD individuals older than 6 years[28, 29]. However, other studies reported that the balance of local and global efficiency between structural and functional networks was reduced in ASD adolescents, and inversely correlated with ASD symptom severity[30].

In the present study, we conducted a source-level assessment of early inefficiencies in brain network topology of young children with ASD. Furthermore, we assessed the clinical relevance of these network alterations by exploring generalizable linked dimensions with behavioral and clinical data. To address these objectives, we measured FC using the ‘EEG source connectivity’ method, which reduces the aforementioned volume conduction and allows networks to be identified directly at the cortical level with optimal spatiotemporal resolution[31–33]. We combined the use of HD-EEG source connectivity method with graph theory-based analysis to characterize and quantify these functional networks at the cortical level. Global and local graph metrics were computed from the identified networks to characterize brain regions (node-wise) and connections (edge-wise) in both young typically developing and ASD children. Finally, we used regularized canonical correlation analysis (rCCA) to assess the relationship among altered cortical network properties and cognitive, sensory and social dimensions within the autism spectrum, to determine whether atypical brain network connectivity may account for individual differences across ASD-related symptomatology and behaviors.

## Materials and Methods

### Participants

This study initially included 50 children diagnosed with autism spectrum disorder (ASD) and 73 typically developing (TD) children. However, 12 children with ASD and 7 TD did not pass the quality check of the HD-EEG data and were excluded from analyses (see below for details), resulting in a final total sample size of 104 participants (**Table 1**). All participants were aged from 2 to 10 years. All children in the ASD cohort were referred to the *Service des Troubles du Spectre de l’Autisme* at Lausanne University Hospital (STSA, CHUV). Trained licensed psychologists and child psychiatrists established formal ASD diagnosis based on the criteria of the Diagnostic and Statistical Manual of Mental Disorders, 5th edition, Text Revision (DSM-5-TR[1]). Clinical procedure for the ASD diagnosis included a review of patients’ medical and developmental history as well as the assessment with the Autism Diagnostic Interview–revised (ADI-R[34]), and the Autism Diagnosis Observation Scale-2 (ADOS-2[35]). TD children were recruited from the general population through distribution of flyers to schools and pediatricians. Exclusion criteria included prematurity (<36 weeks of gestation), a known neurological condition, NDD or learning disability, and/or a first-degree relative diagnosed with ASD. In both cohorts, parents or legal guardians responding to caregiver-report questionnaires were required to be native or fluent French speakers. The study was reviewed and approved by the *Commission Cantonale d’Ethique de la Recherche sur l’être humain* (CER-VD, Switzerland), with the Project-ID 2018-00599. Written informed consents were obtained from participants’ legal representatives before their participation.

**Table 1.**
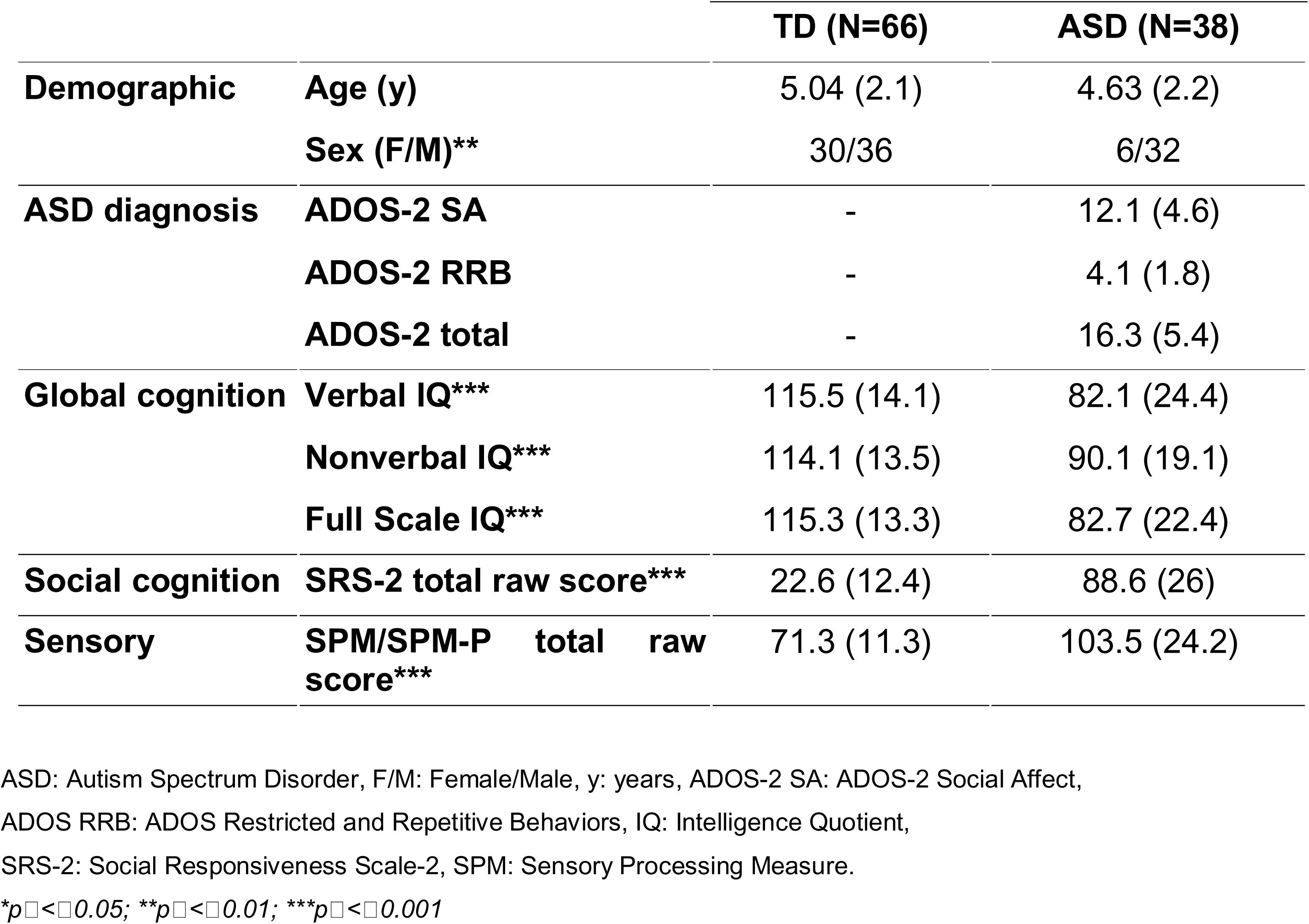
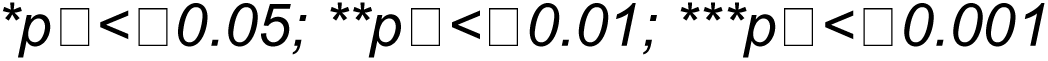
Demographic and clinical characteristics of the study cohort of TD and ASD children expressed as Mean (SD).

|  |  | TD (N=66) | ASD (N=38) |
| --- | --- | --- | --- |
| **Demographic** | **Age (y)** | 5.04 (2.1) | 4.63 (2.2) |
|  | **Sex (F/M)\*\*** | 30/36 | 6/32 |
| **ASD diagnosis** | **ADOS-2 SA** | - | 12.1 (4.6) |
|  | **ADOS-2 RRB** | - | 4.1 (1.8) |
|  | **ADOS-2 total** | - | 16.3 (5.4) |
| **Global cognition** | **Verbal IQ\*\*\*** | 115.5 (14.1) | 82.1 (24.4) |
|  | **Nonverbal IQ\*\*\*** | 114.1 (13.5) | 90.1 (19.1) |
|  | **Full Scale IQ\*\*\*** | 115.3 (13.3) | 82.7 (22.4) |
| **Social cognition** | **SRS-2 total raw score\*\*\*** | 22.6 (12.4) | 88.6 (26) |
| **Sensory** | **SPM/SPM-P total raw score\*\*\*** | 71.3 (11.3) | 103.5 (24.2) |

ASD: Autism Spectrum Disorder, F/M: Female/Male, y: years, ADOS-2 SA: ADOS-2 Social Affect, ADOS RRB: ADOS Restricted and Repetitive Behaviors, IQ: Intelligence Quotient, SRS-2: Social Responsiveness Scale-2, SPM: Sensory Processing Measure.
\* $p < 0.05$ ; \*\* $p < 0.01$ ; \*\*\* $p < 0.001$

### Phenotypic evaluation

#### Overall cognitive functioning

Cognitive ability was measured using standardized tests according to the age and the developmental level of the participant. We used two different tests to measure overall cognitive abilities (intelligence quotient, IQ) in children from 2 years 6 months to 10 years of age: The Wechsler Preschool and Primary Scale of Intelligence (WPPSI-IV)[36] and the Wechsler Intelligence Scale for Children, 5th edition (WISC-V)[37]. Both test batteries included verbal and nonverbal subscales. We used full scale (FSIQ), verbal (VIQ) and nonverbal IQ (NVIQ) standardized to a mean of 100 and a SD of 15, as the outcome measures of cognitive level. For younger children and those not being able to comply with the Wechsler scales, we used the Mullen Scales of Early Learning (MSEL)[38]. We used MSEL Visual Reception subscale as the outcome measure for global nonverbal abilities. In the ASD group, 14 individuals were evaluated with MSEL, 9 with WISC-V and 24 with WPPSI-IV. Finally, 9 TD were assessed with the MSEL, 24 with WISC-V and 33 with WPPSI-IV.

#### Social Responsiveness Scale

Parents completed either the Preschool (2.5–4.5 years old) or the School-age (4.5-18 years old) Social Responsiveness Scale 2nd edition (SRS-2)[39] – a 65-item rating scale measuring deficits in social behavior associated with ASD– about their offspring. In addition to a Total score reflecting a unitary construct of the severity of social deficits, SRS-2 also provides five subscale scores that reportedly constitute treatment subscales including Social Awareness, Social Cognition, Social Communication, Social Motivation, and Mannerisms. We used normally distributed, standardized T-scores, interpreted as: ≤ 59T, within normal limits; 60–65 T, mild; 66–75 T, moderate; ≥ 76T severe[39].

#### Sensory processing questionnaire

The Sensory Processing Measure (SPM)[40] and the Sensory Processing Measure—Preschool (SPM-P)[41] are parent-report questionnaires covering a range of behaviors and characteristics related to sensory processing, social participation and praxis. The age range for SPM-P and SPM was 2–5 and 5–12 years old, respectively. Both SPM-P and SPM include 75 items divided into eight subscales: Social Participation (SOC), Planning and Ideas (PLA), Vision (VIS), Hearing (HEA), Touch (TOU), Taste and Smell (TAS), Body Awareness (BOD) and Balance and Motion (BAL). The last two subscales refer to internal sensory modalities —proprioception and vestibular system—, respectively. A raw total Sensory Score (TOT) was calculated based on the raw score of six sensory system subscales (VIS, HEA, TOU, TAS, BOD and BAL). SOC and PLA refer to higher integrative functions (social functioning and praxis, respectively) and hence did not contribute to the composite score. Standardized T-scores for each subscale except for TAS were used as outcome measures. SPM was used for a sample of 16 ASD and 33 TD children. Twenty-two ASD individuals and 33 TD were assessed with SPM-P version.

### Electrophysiological acquisition and preprocessing

#### Acquisition

HD-EEG data were collected using a 128-channel HydroCel geodesic sensor net together with Electrical Geodesics, Inc (EGI, Eugene, Oregon) Net Station software[42]. This system uses the vertex (Cz) as physical reference. Prior to place the cap on the child’s head, the net was soaked in an electrolyte solution to facilitate electrical contact between the scalp and the electrodes. Individual sensors were adjusted until impedances were kept below 50 kΩ. HD-EEG signals were sampled at an effective rate of 250 Hz. All channel signals were amplified using NetAmps 200 amplifier and digitized with a 12-bit A/D converter. Resting-state HD-EEG was collected during 5 minutes in a dark room as children watched an age-appropriate soundless video presented in a computer screen while seated on a chair or on their caregiver’s lap. A research assistant accompanied the child to keep motivation and cooperation during HD-EEG data acquisition.

#### Preprocessing

Raw HD-EEG signals were bandpass filtered off-line in the range 0.1-40Hz. Filtered HD-EEG data was further segmented into epochs of 40 seconds each, and the first segmented epoch was always discarded for later analyses. HD-EEG epochs were preprocessed with MATLAB R2020b software (MathWorks) and the open-source toolbox preprocessing pipeline Automagic (v2.5)[43]. This toolbox simplifies and monitors artifacts removal, interpolation of error-prone channels and data quality rating (**Figure 1A**). The remaining artifact-free epochs were sorted based on the overall high-amplitude, and only the first three epochs of 40 sec were selected for analysis. A trained EEG expert (AM) reviewed the decisions made by Automagic, and the epoch selection process underwent confirmation through visual inspection, with any epochs displaying lingering artifacts subject to modification or complete removal. Over 84% of the HD-EEG data exhibited ‘ok’ or ‘good’ data quality. As a result, a total of 12 children with ASD and 7 TD children exhibited HD-EEG signals categorized as ‘bad’ and thus met criteria for exclusion of later analyses, leaving the final sample for analysis in 38 ASD and 66 TD children.

**Figure 1.**
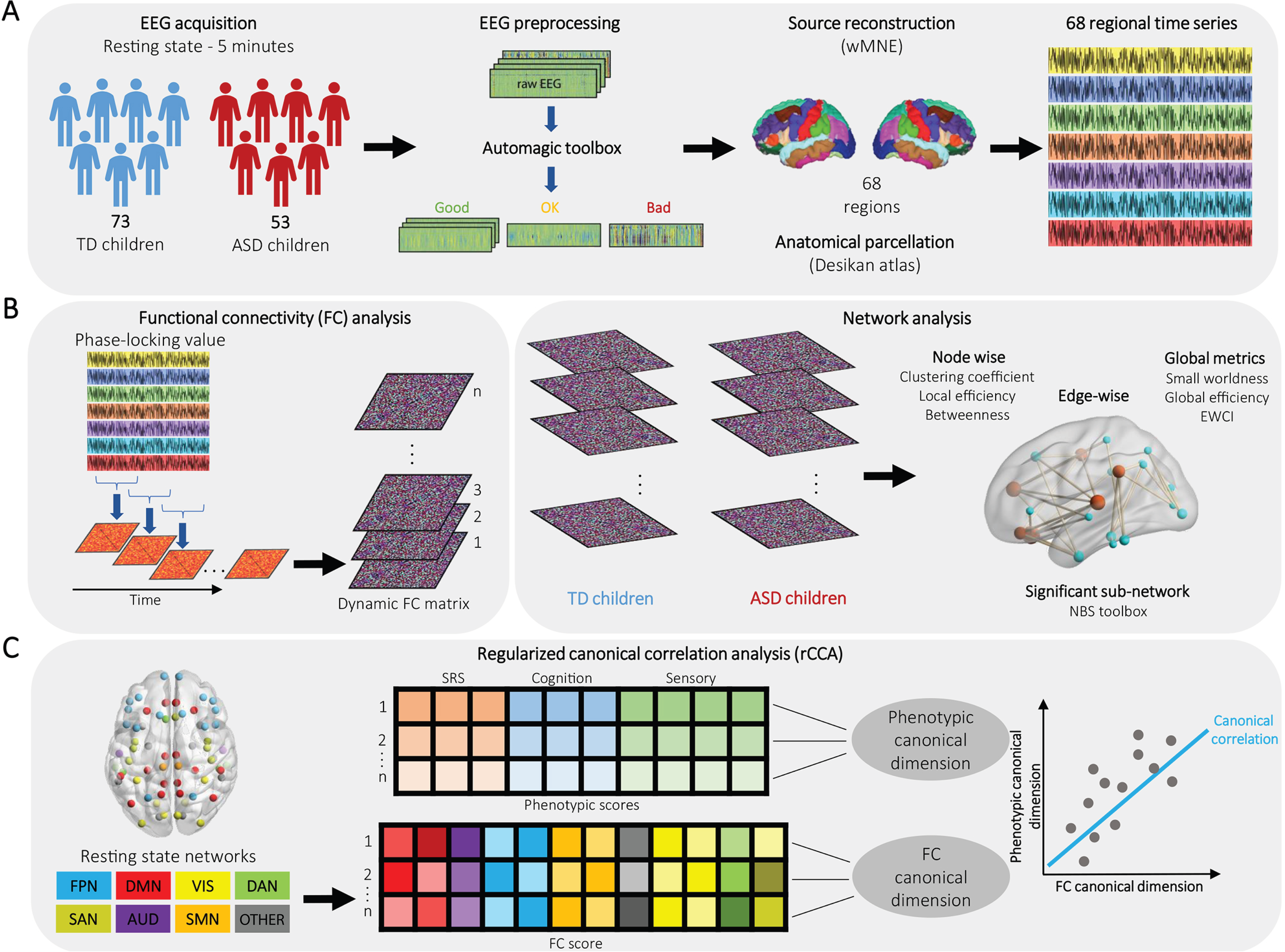
**A.** Schematics summarizing pre-processing pipeline and cortical parcellation. Automagic toolbox was used for quality assessment and artifact correction, and the cortical sources were reconstructed by solving the inverse problem. An anatomical parcellation was applied on the adapted MRI template yielding a time-course RS HD-EEG signal for 68 cortical regions of interest using the Desikan-Killany atlas. **B.** FC between the 68 regional time series was computed for each time window using the Phase Locking Value (PLV) method. A resulting dynamic FC matrix per individual was estimated by the standard deviation across all sliding windows. We compared dynamic FC matrices between ASD and TD young children using three levels of network analysis: i) *node-wise*: we computed metrics such as clustering coefficient (CC), local efficiency (LE), and betweenness centrality (BC); ii) *edge-wise*: we retrieved the cortical subnetwork whose edges underwent a significant change in FC values after statistical comparison of ASD and TD connectivity matrices using NBS approach; and iii) *global network topology*: Small World Propensity (SWP), edge-wise connectivity index (EWCI) and global efficiency (GE) were calculated to reflect overall network organization. **C.** Regularized canonical correlation analysis (rCCA). The 68 parcellated cortical regions were grouped into eight large-scale network communities as described in Suppl. Figure 1. FC features for each of these network communities as well as cognitive, social and sensory phenotypic scores were entered into rCCA to search for a canonical pair that maximizes the correlation between FC and behavioral dimensions.

#### Source reconstruction

To obtain estimates of brain activity at the cortical level, we applied the first step of the EEG source connectivity method[44]. We reconstructed the dynamics of cortical brain sources from the artifact-free HD-EEG data segments by solving the inverse problem. An atlas-based approach was used to project HD-EEG sensor signals onto an anatomical framework consisting of 68 cortical regions identified by means of the Desikan-Killiany cortical atlas[45] using Freesurfer[46]. We built an age-matched 4-to-8 years head model using the *NIH* Pediatric *MRI* database (http://www.nitrc.org/projects/pediatric_mri/).

Using the OpenMEEG toolbox[47], the procedure included the co-registration of the pre-processed HD-EEG signal with the MRI template through identification of the right/left pre-auricular points and nasion anatomical landmarks. The lead field matrix was then computed for a cortical mesh with 15,000 vertices using Brainstorm[48]. The weighted minimum norm estimate (wMNE)[49] was used to calculate the temporal dynamics of the 68 cortical regions of interest (ROIs).

#### Functional connectivity (FC)

The second step of the EEG source connectivity method involved the estimation of the statistical couplings between the 68 source-reconstructed regional time series separately for four different frequency bands: theta (4–8 Hz), alpha (8– 13 Hz), beta (13–30 Hz) and gamma (30-45 Hz). FC between the regional time series was computed for each frequency band using the Phase locking value (PLV) method[50]. This measure, ranging between 0 and 1, reflects interactions between two oscillatory signals through quantification of the phase relationships. The combination of the wMNE and the PLV methods was chosen according to a recent model-based comparative study of different inverse/connectivity combinations[51]. We obtained for each participant four dynamic FC matrices in each frequency band. Those matrices were ultimately averaged across time and epochs to obtain a single static FC matrix per frequency band, used in the further analysis. This resulted in a 68×68 symmetrical FC matrix from which 68×67/2= 2,278 unique connections were obtained in each frequency band, representing the pairwise connections between all of the cortical ROIs (**Figure 1B**).

#### Network metrics

FC matrices were characterized using network measures derived from graph theory, in a system in which nodes (68 cortical regions) are connected by edges reflecting the functional and/or effective connections. Then, network topological properties were quantified to characterize brain network architecture using the Brain Connectivity Toolbox (BCT)[52] and visualized with BrainNet viewer[53]. Weighted undirected networks were constructed in which the connection strength between each pair of nodes was defined as their connectivity value. Functional networks were analyzed at three different levels: brain regions (node-wise), connections (edge-wise) and global (the whole brain). The equations to compute all network metrics described below are detailed in **Supplementary Methods**.

##### Node-wise

We measured clustering coefficient (CC), betweenness centrality (BC) and local efficiency (LC). CC is a major metric for determining the level of information segregation of the network, by showing how many nodes are clustered together in a graph (i.e., the fraction of node’s neighbors that are neighbors of each other)[52]. The more the neighborhood of the given node is densely interconnected, the higher is its local CC. BC of a node indicates the importance and the amount of information flowing via that node[54]. To compute BC, we first calculated the shortest path length (SPL) between nodes, that is, the minimum number of links to be traversed in the optimal path from one node to the other. BC is the fraction of all shortest paths in the network that contain a given node. Nodes with high BC values participate in a large number of shortest paths. LE is an average measure of efficiency of information transfer limited to neighboring nodes, defined as the average inverse SPL of all neighbors of a given node among themselves[55].

##### Edge-wise

Network metrics were characterized using the Network-Based Statistic (NBS) toolbox[56]. To do that, an ANCOVA analysis was fitted to each of the 2,278 PS values of each of the edges, yielding a p-value indicating the probability of rejecting the null hypothesis at each edge. A component-forming threshold was applied to each p-value, and we obtained the size of each connected element in these thresholded matrices. The size of the components was then compared to a null distribution of maximal component sizes obtained using permutation testing to obtain p-values corrected for multiple comparisons[56]. The NBS method finds subnetworks of connections significantly larger than would be expected by chance. Here we report results for a threshold that retain only edges with p < 0.005.

##### Global topology

Small World Propensity (SWP)[57] has been proposed as a reliable measure to estimate information transfer in a network. SWP evaluates the level of small-world structure between any two nodes, that is, higher local clustering than a random network, yet short average path length. Furthermore, we measured global efficiency (GE) to further evaluate differences in the integration of the neural networks of children with ASD and TD children. GE measures the efficiency of information transfer among all pairs of nodes in the graph, and thus reflects the efficiency of interaction across the whole graph[52, 58]. When applying NBS to compare the networks of the two groups, we computed an Edge-wise Connectivity Index (EWCI), defined as the sum of the weights of the resultant subnetwork[59].

#### Statistical analysis

Shapiro-Wilk normality tests were conducted to determine the normality distributions of the network metrics. When normality was not met, Wilcoxon signed-rank tests were used. Group differences of power spectral density and network metrics between ASD and TD children were assessed with an Analysis of Covariance (ANCOVA) model in R (v4.2.1)[60]. We used the group factor (ASD or TD) as the main variable, and age and sex as covariates. Levene’s test was used to assess equality of variance. We also assessed the developmental trajectory of global network metrics as a function of age in the TD and ASD groups with a regression model including linear and quadratic expansions of age. Group differences in developmental trajectory were modeled with the network metric (GE, SWP, EWCI) as the dependent variable and group, age, age^2^ and groupXage interaction independent variables. We performed a quantitative node-wise analysis on the brain networks of TD and ASD children using BC, CC and LE as network metrics. P-values for node features were corrected for multiple comparisons using FDR across the 68 nodes for each metric. To further explore how edge-wise differences might manifest in pairwise –ROI to ROI– connectivity strengths, we tested for FC differences in each of the 2’278 possible connections. P-values were corrected by permutation test using NBS. Effect sizes are reported as well in the form of Cohen’s D for two-sided Welch’s t-tests, and standardized beta values for regressions. We generated Cohen’s d maps from FDR-corrected t scores to show the unbiased magnitude of the effects.

### Regularized canonical correlation analysis (rCCA)

Canonical correlation analysis (CCA) is a multivariate approach to highlight correlations between two data sets. A regularized modification of canonical correlation analysis (rCCA) which imposes an L2 penalty (λ) on the CCA coefficients is widely used in applications with high-dimensional data and/or those with high collinearities[61, 62]. We used rCCA to identify latent generalizable brain–behavior linked dimensions explaining individual differences in three ASD-related domains: social, cognitive and sensory processing behaviors (**Figure 1C**). We performed three L2-regularized CCA using the rCCA function in R mixOmics package[63], separately for the three abovementioned clinical domains.

To avoid potential overfitting and handle multicollinearity in the input variable sets, we implemented a robust selection of clinically meaningful FC features and define a low-dimensional representation for those FC features, as detailed in **Supplementary Methods**. Moreover, we regressed age and sex out of both the FC and the phenotypic data to limit the influence of potential confounding factors. To help understand the cortical networks that are highly contributing to the identified FC canonical dimensions, the ranked FC loadings were parcellated and assigned to one of the eight well-established large-scale cortical within- and between-network communities (**Suppl. Figure S1; Suppl. Table S1**)[64]: visual network (VIS), auditory network (AUD), sensorimotor network (SMN), fronto-parietal network (FPN), salience network (SAN), dorsal attentional network (DAN), default mode network (DMN) and other networks (OTHER). FC loadings assigned to each network community were summed as measures of network strength[65, 66], establishing the final number of FC features incorporated to the rCCA in (8×8/2=36). We generated two canonical variate pairs (FC and behavior scores) with values for all participants and a canonical correlation for each variate pair. To better illustrate the features that stably contribute to each canonical dimension (FC and behavior), we computed regression coefficients (i.e. canonical loadings) between each canonical variate and its original variables[67]. Loadings represent the weights assigned to each of the original variables to determine their contribution to a given latent dimension. The canonical loadings are crucial for the interpretation step as higher loading values indicate greater contribution of the given variable to the canonical dimension. We reported canonical dimensions that were statistically significant at false discovery rate correction (FDR, q < 0.05) and which explain more than 5% of the covariance for further analyses[68].

## Results

### Demographics and clinical characteristics

Demographics and clinical scores are presented in **Table 1**. Age was not significantly different between TD and ASD children (p===0.35), but we observed a higher prevalence of males in the ASD group (χ^2^ = 8.1, p = 0.005). ASD children had a significantly lower global cognitive functioning compared to TDs, with an average of 33 points lower VIQ (p===6.3e-10) and a 24-point lower NVIQ (p===8.3e−9). ASD children scored higher in both the SRS-2 (p===1.02e−17) and the SPM total score (p===1.26e−9). SRS-2 and SPM subscales scores are detailed in **Supplementary Table S2**.

### Frequency-based analysis

To validate the consistency of our EEG data spectral power, we analyzed the overall changes in the relative spectral power of ASD and TD children over all brain regions (**Figure 2A**). We confirmed that the peak power fell within the alpha band for both groups, as expected in task-free EEG recordings[69]. Higher relative power seems to characterize the neural oscillatory activity of ASD children at lower frequency bands. **Figure 2B** summarizes the frequency-based analysis, showing an increase in the power spectral density of ASD children in the delta and alpha frequency bands, although only the differences in the alpha band survived correction for multiple comparisons (p < 0.01, Bonferroni corrected). We found no significant differences in neither theta, beta nor gamma frequency bands. The results presented in the following sections are then circumscribed to alpha band (8-13 Hz).

**Figure 2.**
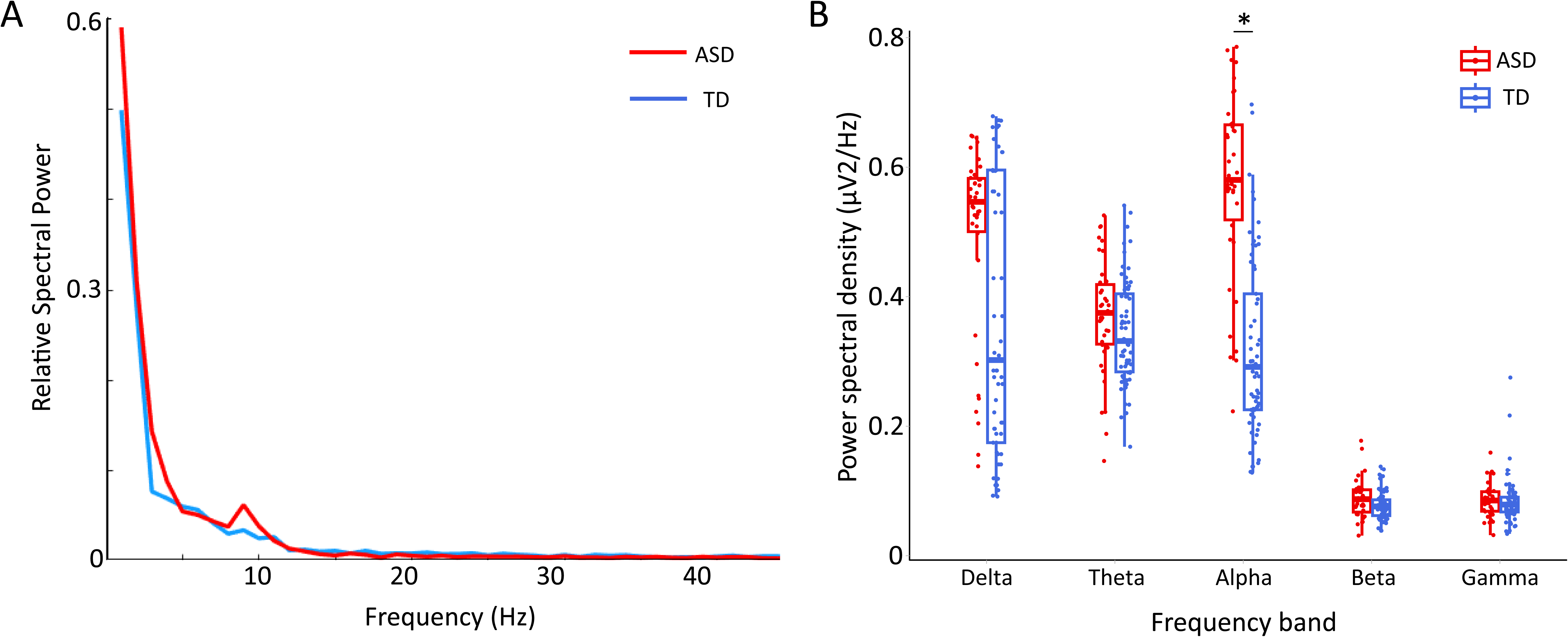
**A.** Power spectral representation of EEG data for ASD (red) and TD (blue) young children. **B.** Frequency-based analysis: Individual values of the power spectral density for ASD and TD young children at five frequency bands: [delta (0.5–4 Hz); theta (4–8 Hz); alpha (8–13 Hz); beta (13–30 Hz); gamma (30– 45 Hz)]. In all boxplots, the thick horizontal line shows the median, and the bottom and top of the box show the 25th (quartile 1 [Q1]) and the 75th (quartile 3 [Q3]) percentile, respectively. The upper whisker ends at highest observed data value within the span from Q3 to Q3ϑ+ϑ1.5 times the interquartile range (Q3–Q1), and lower whisker ends at lowest observed data value within the span for Q1 to Q1-(1.5 * interquartile range). Points not reached by the whiskers are outliers. Significant post-hoc group comparisons are represented by solid lines above. *pϑ<ϑ.01, Bonferroni-corrected.

### Mapping of cortical topological differences

#### Node-wise

Differences in node-wise network metrics between ASD and TD children are displayed in **Figure 3**. The results highlight higher CC in the network nodes corresponding to frontal areas –the right lateral, medial and superior orbitofrontal cortex (d=-0.43, p=0.011)–, as well as the lateral occipital and pericalcarine cortex of the left occipital lobe (d=-0.66, p=1.6e-4) of ASD children. These results present a distinct symmetry between the two hemispheres. We also found that ASD children exhibited higher CC in the precuneus, a part of the superior parietal lobule, of the right hemisphere (d=-0.25, p=0.03). Nevertheless, only the higher CC observed in the left occipital lobe of ASD children persisted after FDR-correction. Similarly, we found the largest differences in LE also in the left lateral occipital region (d=-0.67, p=1.4e-4). Other regions such as the right entorhinal and middle frontal cortices, as well as the left superior temporal gyrus, presented trends for a higher LE in ASD children, but they did not survive correction for multiple comparisons. Lastly, we found significantly higher BC in both the right inferior parietal and rostral part of the inferior frontal gyrus of ASD children (d=-0.37, p=2.6e-4). Thus, network nodes with significantly different BC do not seem to be symmetrically distributed across the hemispheres but involve predominantly the right one. Smaller trends in the left parsopercularis and superior temporal gyrus did not persist after FDR-correction.

**Figure 3.**
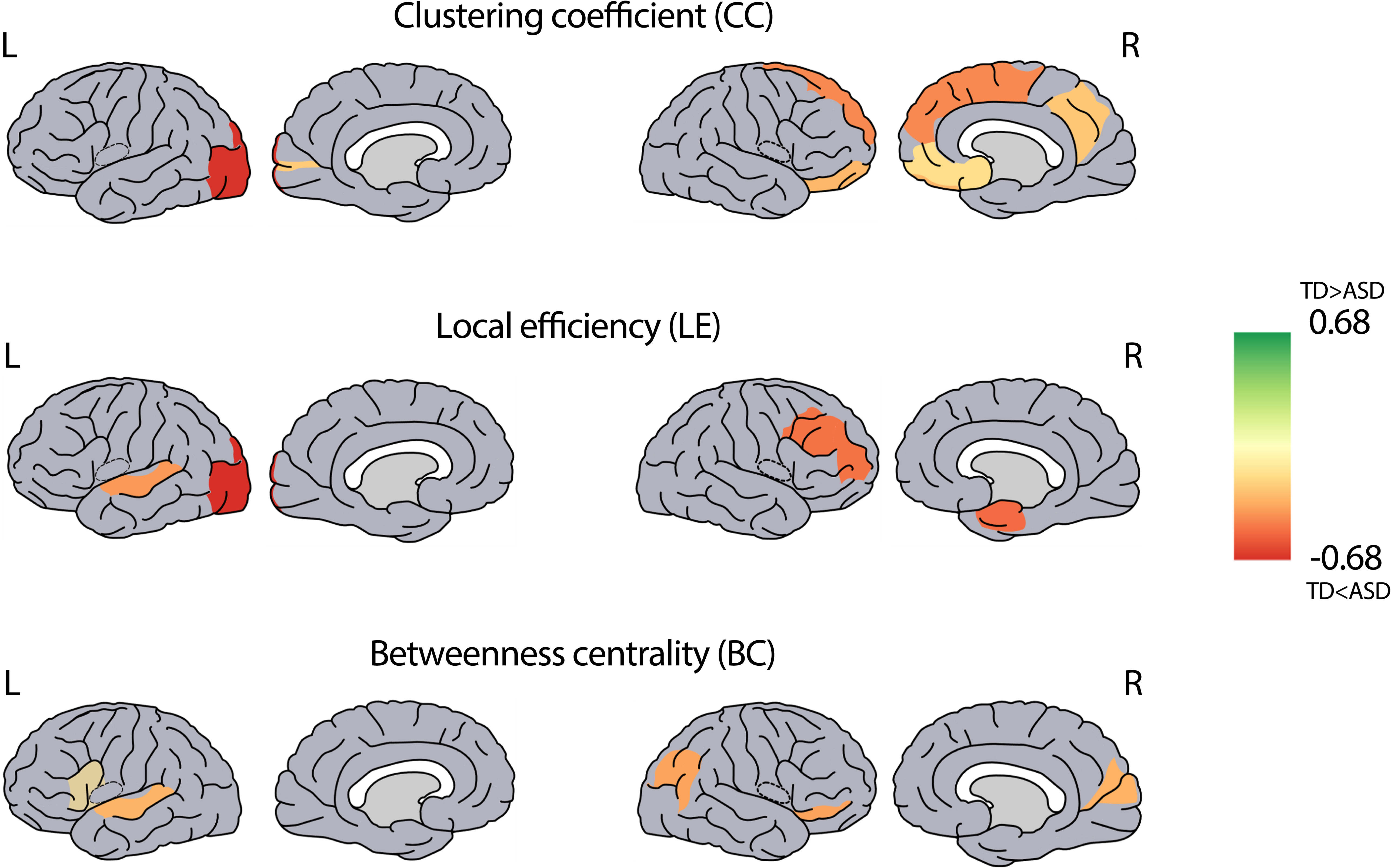
Node-wise analysis. Cortical mapping of local network overconnectivity in young children with ASD. Cohen’s d maps from FDR-corrected t scores in clustering coefficient (upper), local efficiency (middle) and betweenness centrality (lower) are displayed. Color bars represent Cohen’s d. *L, left; R, right.* The spatial distributions of significant brain regions which is represented as a sphere. A-.

**Table 2.** Node-wise results: List of cortical regions with significantly different network metrics between TD and ASD children expressed as Mean (SD).

|  |  | TD | ASD | Cohen D | p-value |
| --- | --- | --- | --- | --- | --- |
| **CC** | **R lateral orbitofrontal** | 0.49 (0.02) | 0.62 (0.07) | -0.31 | 0.02 |
|  | **R medial orbitofrontal** | 0.35 (0.02) | 0.77 (0.07) | -0.17 | 0.01 |
|  | **R superior frontal** | 0.49 (0.02) | 0.62 (0.06) | -0.43 | 0.03 |
|  | **R precuneus** | 0.48 (0.03) | 0.65 (0.08) | -0.25 | 0.02 |
|  | **L pericalcarine** | 0.51 (0.03) | 0.63 (0.08) | -0.24 | 0.03 |
|  | **L lateral occipital** | 0.17 (0.03) | 0.59 (0.05) | -0.67 | **1.5e-4** |
| **LE** | **R rostral middlefrontal** | 0.4 (0.18) | 0.57 (0.07) | -0.49 | 0.01 |
|  | **R entorhinal** | 0.42 (0.31) | 0.62 (0.14) | -0.43 | 0.01 |
|  | **L superior temporal** | 0.49 (0.17) | 0.61 (0.07) | -0.48 | 0.02 |
|  | **L lateral occipital** | 0.135 (0.02) | 0.59 (0.04) | -0.68 | **1.4e-4** |
| **BC** | **R inferior parietal** | 0.05 (0.001) | 5.31 (0.32) | -0.36 | **2.6e-4** |
|  | **R cuneus** | 0.09 (0.002) | 2.825 (0.27) | -0.33 | 0.01 |
|  | **R pars orbitalis** | 0.03 (0.001) | 1.815 (0.2) | -0.37 | 0.01 |
|  | **L superior temporal** | 0.087 (0.001) | 0.712 (0.01) | -0.32 | 0.02 |
|  | **L pars opercularis** | 0.03 (0.005) | 0.261 (0.09) | -0.1 | 0.03 |

#### Edge-wise

The results of the edge-wise analysis are summarized in **Figure 4**. The network reconfiguration in the alpha band that emerges when comparing ASD and TD children, as evaluated by the change in FC values, comprises 30 edges between 22 brain regions (p<0.05 after correction for multiple comparisons). All these edges exhibited significantly increased FC in ASD children. Spatially, the resulting network is predominantly localized in the anterior areas of the brain, presenting connections across the hemispheres, as well as intra-hemisphere connections on the right anterior and left posterior sides (**Figure 4A**). The significant connections are predominantly frontotemporal, involving mainly the right frontal (lateral and medial orbitofrontal, as well as rostral middlefrontal), and the left temporal (transverse temporal gyrus) lobes. These results are in accordance with the nodewise analysis results reported in the previous section. To better characterize the mapping of the edges, we calculate the degree for each of the 22 brain regions within the resulting network, which represents the number of connections (edges) associated to each specific node (**Figure 4B**). We confirmed that the right lateral orbitofrontal cortex is the region with highest connection density (degree=6) in the resulting network with significantly increased FC in ASD children. Other highly interconnected regions in this network are the left transverse temporal and the right rostral middle gyri.

**Figure 4.**
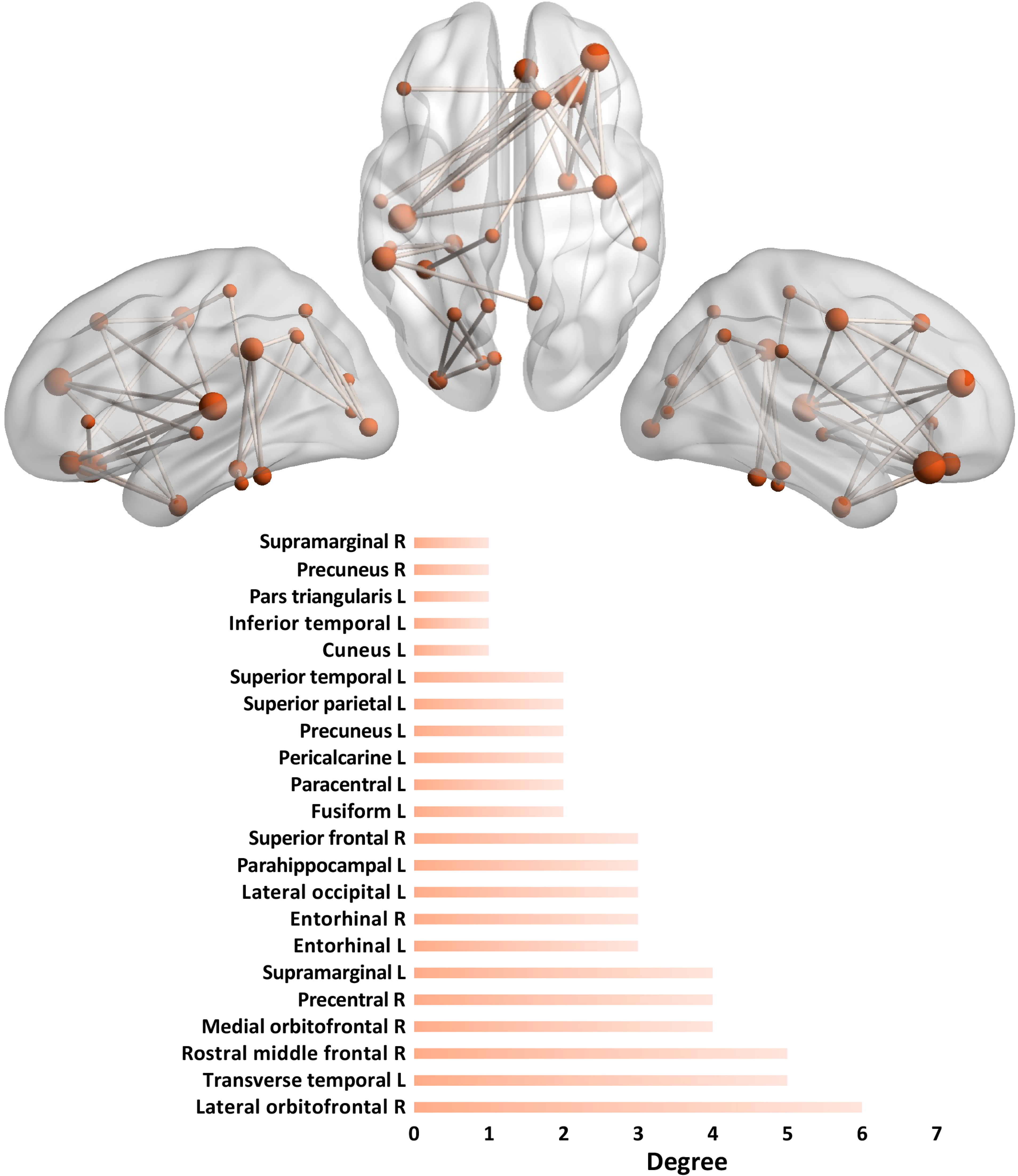
Edge-wise analysis. Graphical representation (upper)of the resulting network showing the 30 FDR-corrected edges with higher FC in young children with ASD. Each cortical region (node) is represented as a sphere plotted according to the stereotactic coordinates of its centroid, and each suprathreshold edge represents the connections between the regions as a solid line. The size of the node is proportional to its degree, i.e. the number of connections in which it takes part. In the lower panel, representation of the degree for each brain region identified in the resulting network. P-values were corrected by permutation test using NBS (p < 0.05). *L, left; R, right*.

#### Group differences and developmental trajectories of global network metrics

Global estimates and developmental trajectories of the analyzed network metrics are presented in **Figure 5**. The group comparison showed that SWP was comparable between ASD and TD children (t_(103)_===0.13, p===0.89, **Figure 5A**). We computed the EWCI using the resulting network from the edge-wise analysis consisting of 30 edges and 22 brain regions. EWCI was significantly higher in ASD compared to TD children (t_(103)_===3.21, p===0.002, **Figure 5B**). Lastly, GE presented a trend to be lower in ASD children (t_(103)_===-2.06, p===0.047), but did not survive FDR correction (**Figure 5C**). The analysis of the developmental trajectory of these network metrics revealed that the increased EWCI was steadily constant and already present at an early age. We also observed that SWP did not significantly change with age in TD children (r^2^ = 0.01, p===0.71), whereas ASD children exhibited a U-shaped quadratic trajectory with higher SWP at specifically older ages (r^2^ = 0.26, p===0.004). Finally, GE showed no change over time in both ASD (r^2^ = 0.06, p===0.16) and TD (r^2^ = 0.03, p===0.61) groups.

**Figure 5.**
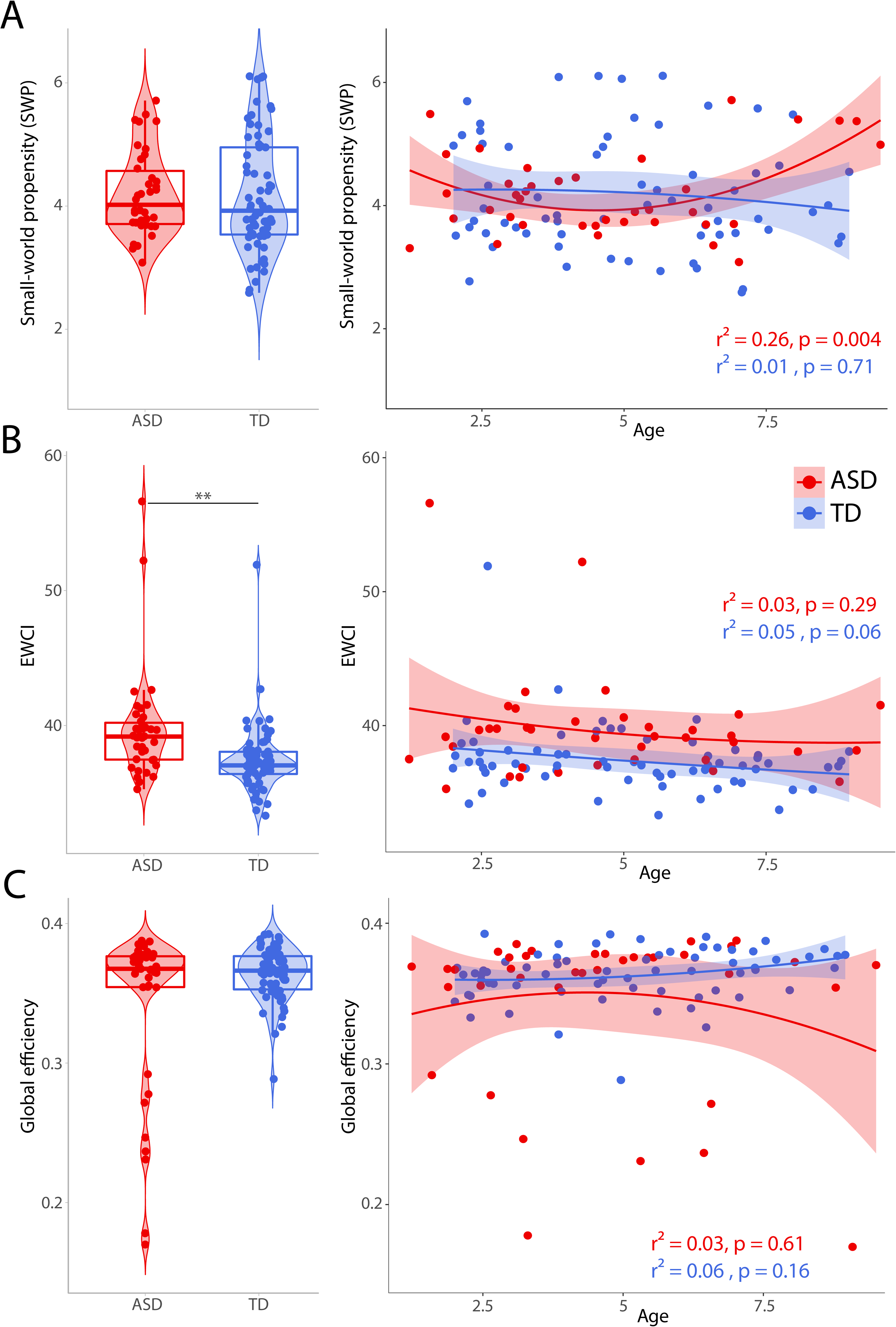
Global network topology. For each metric (**A**, small-world propensity; **B**, EWCI; **C**, global efficiency) the left panel shows group comparison of individual values between ASD and TD young children, whereas the right panel displays age-related changes in global network topology for both groups. EWCI was computed from the resulting network of the edge-wise analysis. Linear and quadratic (2^nd^ order) regressions of age were modeled for each global network metric. In all boxplots, the thick horizontal line shows the median, and the bottom and top of the box show the 25th (quartile 1 [Q1]) and the 75th (quartile 3 [Q3]) percentile, respectively. The upper whisker ends at highest observed data value within the span from Q3 to Q3ϑ+ϑ1.5 times the interquartile range (Q3–Q1), and lower whisker ends at lowest observed data value within the span for Q1 to Q1-(1.5 * interquartile range). Points not reached by the whiskers are outliers. **pϑ<ϑ.01, FDR-corrected.

### Linked brain-behavior dimensions

We conducted three rCCA as a data-driven approach to unfold linked dimensions between brain FC and social, cognitive and sensory processing features, respectively, in both ASD and TD children. Following the grid search of optimized regularization parameters of rCCA (**Supplementary Fig. S2**), we selected for further analysis the canonical pairs based on the accumulated covariance explained. For the FC-cognition rCCA, the most significant canonical variate pairs (r^2^ = 0.44, p_FDR_ = 4.5e-6) explained 67% of variance of the cognitive features, and 17% of variance of FC features, respectively (**Figure 6A**). Based on their highly weighted item compositions, the cognitive canonical dimension was characterized by higher loadings of nonverbal skills. At the network level, this cognitive dimension was mainly associated to overconnectivity in DAN-SMN (network strength = 0.76) and DAN-VIS (network strength = 0.73) between-network pairs. The FC-sensory rCCA revealed a canonical pair (r^2^ = 0.63, p_FDR_ = 3.4e-12) explaining 62.8% of sensory processing variance, as well as 11.3% of FC variance (**Figure 6B**). Auditory, tactile and proprioception were the most contributing modalities to the sensory canonical variate. This dimension was mainly correlated with higher FC in ASD children between DAN-SMN networks (network strength = 1.58). Finally, the most powerful canonical pair (r^2^ = 0.54, p_FDR_ = 5.8e-9) yielded by the FC-Social rCCA explained 84.7% and 10.7% of SRS and FC variance, respectively (**Figure 6C**). The social canonical dimension was characterized by high loadings of social motivation and restricted and repetitive behavior subscales. ASD children exhibited higher values in this social dimension that were associated with increased FC prominently between DAN-FPN (network strength = 1.24) and DAN-SMN (network strength = 1.17) networks.

**Figure 6.**
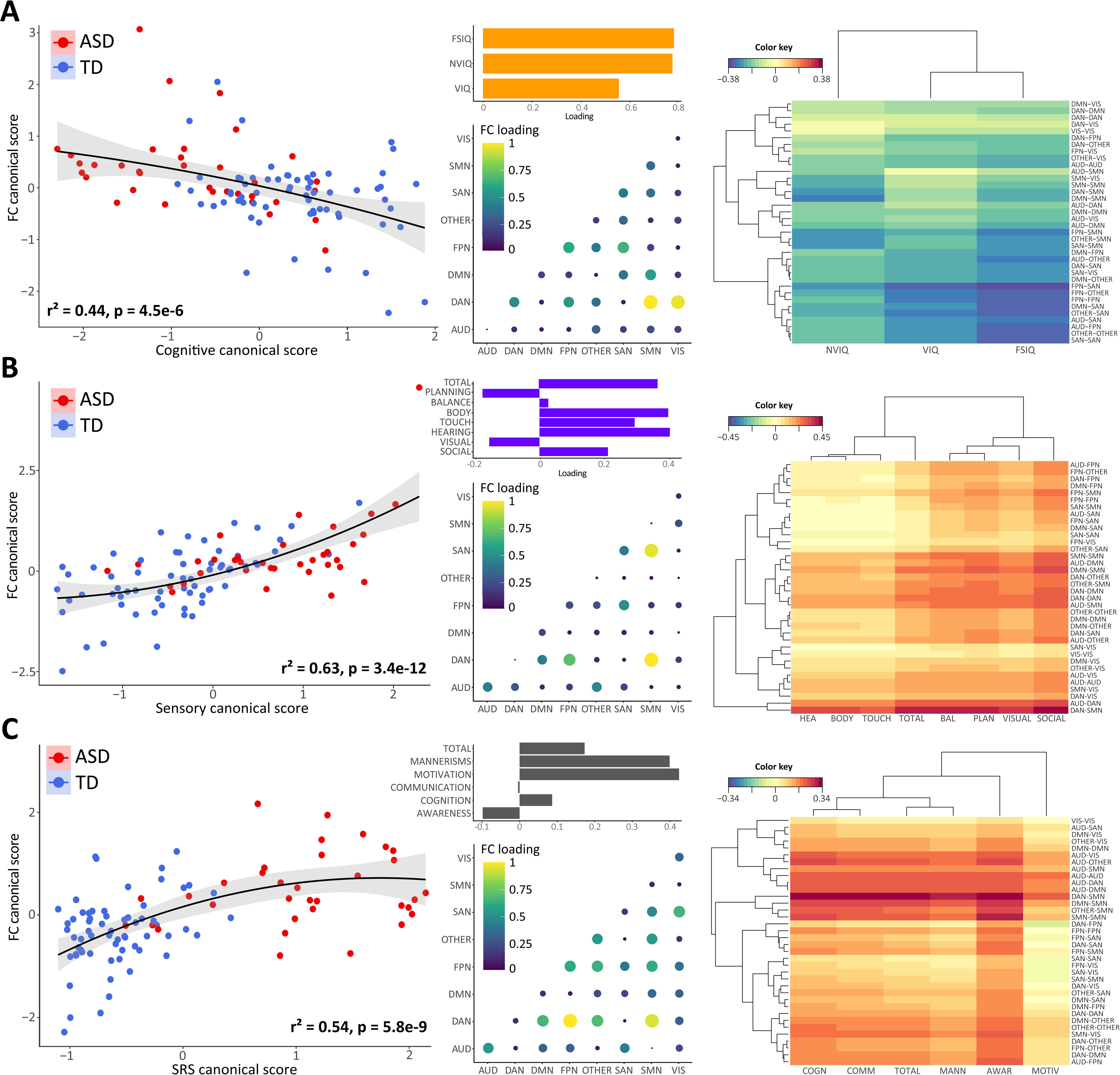
Linked brain–behavior dimensions to predict individual differences in cognitive (**A**), social (**B**) and sensory processing (**C**) scores using network connectivity patterns in young ASD and TD children. Left panels are scatterplots depicting linear correlations of FC with behavioral scores: cognitive (r^2^ = 0.44), social (r^2^ = 0.63) and sensory processing (r^2^ = 0.54). The middle panel shows the loadings of behavioral and FC features, separately for each brain and behavior dimension. On the right, Clustered Image Maps representing Pearson correlation coefficients between FC and behavior datasets. Hierarchical clustering is applied on the FC (rows) and behavior (columns) features simultaneously. Dendrograms are used along the axes to depict how each row/column clusters based on the hierarchical clustering method.

## Discussion

In the present study, we used source-level reconstruction from resting-state HD-EEG to characterize functional network overconnectivity, and its association with cognitive, sensory and social dimensions in young children with ASD. Compared to a group of TD children, cortical network metrics in the alpha band of ASD children revealed local overconnectivity in frontotemporal and lateraloccipital regions. We unveiled a frontotemporal cross-hemispheric neural connectivity pattern that stably contributed to the FC differences between ASD and typical brain development already from early childhood. Finally, we identified three latent FC– behavior dimensions explaining individual differences in cognitive, social and sensory phenotypic features. Although each linked brain–behavior dimension described a distinct FC pattern, overconnectivity between DAN and SMN networks was a recurrent functional inefficiency contributing to explain variance in the aforementioned ASD-related behavioral dimensions. Taken together, these findings define discrete patterns of cortical functional overconnectivity that help to account for individual differences in cognitive, social and sensory ASD-related manifestations.

Given the acknowledged limitations of univariate methods to establish generalizable brain– behavior dimensions[70], data-driven multivariate approaches provide a reasonable alternative to explore solid relationships between brain features and behavior. In this study, we conducted rCCA to build up maximal correlations between linear combinations of both behavioral and FC data, to better understand how atypical network overconnectivity underlies individual differences in ASD-related phenotypic manifestations. Regularized CCA is a multivariate modeling and predictive analysis that has been validated in biopsychological studies[71, 72]. Assuming that the most highly weighted features for each canonical variate provide a fundamental estimation of its essence, we found that each cognitive, social and sensory brain–behavior dimension described a unique FC pattern. The cortical network overconnectivity most strongly correlated to individual variability in the cognitive dimension was between DAN and SMN networks, as well as between DAN and VIS networks. These results are in agreement with previous resting-state fMRI evidence of a robust association between higher intelligence scores and lower FC in connections between cortical areas comprising DMN and DAN, including the visual, and somatosensory regions[73, 74]. FC patterns contributing to variance in the dimension based on social responsiveness involved increased functional interactions mainly between DAN, FPN and SMN networks. It is important to note that similar patterns of FC-behavior relationships have already been observed in ASD individuals, with higher SRS scores associated with hyperactivity in several DAN and FPN regions relevant for socio-emotional processing, such as the medial frontal and bilateral middle temporal gyrus[75]. There is compelling evidence indicative of reduced segregation of social brain networks in ASD, manifested as an excessive number of connections with irrelevant regions[14]. Finally, greater auditory, tactile and proprioceptive processing difficulties in ASD young children were associated with higher FC between DAN and SMN networks. These results are in line with previous work that demonstrates increased FC of primary sensory networks in ASD individuals[76]. Other studies have established robust relationships between sensory hyper-responsiveness and FC in salience networks[77], but we could not replicate this result with our sample. Recurrent overconnectivity between DAN and SMN networks was shared between all identified brain-behavior dimensions, consistent with previous work showing hyper-connectivity in both dorsal and ventral attentional networks of young ASD children[78]. Our findings thus suggest that precise patterns of network overconnectivity contribute to specific phenotypic manifestations in young children with ASD.

A prevailing theory of developmental FC changes in ASD posits that young ASD children exhibit local overconnectivity, in contrast to underconnectivity in adults with ASD[79, 80]. Here we expand upon previous findings of increased FC in ASD early childhood by characterizing higher-level network properties using tools derived from the physics of complex networks[52]. We report local network overconnectivity in community organization within frontotemporal cross-hemispheric networks and in lateral-occipital regions, in young children with ASD relative to their TD counterparts. It is hypothesized that FC within frontal lobe of ASD individuals is excessive, with disorganized and inadequately selective network properties, whereas FC with other lobes is suboptimal, with reduced responsiveness and deficient transfer of information[81, 82]. We also observed that frontotemporal networks in young children with ASD had shorter average path lengths (i.e., higher levels of local efficiency). Randomly connected networks tend to have shorter path lengths[83], suggesting the possibility that higher efficiency may simply reflect a less organized or more random distribution of functional edges. This is consistent with a study finding decreased complexity or increased randomness in resting-state fMRI timeseries of individuals with ASD[84]. Our node-wise analysis also revealed increased regional segregation in the lateraloccipital and inferior parietal regions. Similar patterns have been observed with other neuroimaging techniques[85–87], suggesting region-specific overconnectivity partially associated with more integrative long-distance FC. Furthermore, local overconnectivity in occipital and posterior temporal regions of children with ASD has been reported, together with underconnectivity in posterior cingulate and medial prefrontal regions[88].

We found higher relative power in the alpha band in young children with ASD. At rest, the EEG signal is characterized by oscillations in the alpha frequency band, which have been associated with attention and perceptual processing tasks[89]. Neural activity in the alpha band is of interest since it is thought to denote the level of cortical excitability[90]. The neural oscillatory pattern exhibited by ASD children in our study seems which is consistent with previous studies[91, 92], and has been associated with cortical deactivation or inactivity[90, 93]. Age-dependent alpha FC patterns have been observed, demonstrating hyperactivation in the ASD group for ages below 6–7 years old and hypoactivation for older ages[27]. However, there is also contrasting evidence suggesting comparable alpha spectral density between children with ASD and TD children[94]. The potential sources of these discrepancies may include differences in the experimental protocols and methodology. For instance, higher levels of posterior alpha activity have been observed in ASD individuals during eyes-open resting-state EEG, seemingly attributable to reduced occipital alpha suppression[95].

Classically, FC has been usually computed at the scalp level, not allowing interpretation of anatomically interacting brain areas as they are severely corrupted by the volume conduction effects[96]. Here, EEG source connectivity method was used to identify functional networks at the cortical level from HD-EEG recordings[31]. As compared with fMRI studies, a unique advantage of this method is that networks could be directly identified at the cerebral cortex level from scalp EEG recordings, which consist in direct measurement of neuronal activity, in contrast with blood-oxygen-level-dependent (BOLD) signals. This method was first evaluated for its capacity to reveal relevant networks in a picture naming task[31], and was then extended to the tracking of the spatiotemporal dynamics of reconstructed brain networks[97]. HD-EEG is a non-invasive, portable, widely available, and cost-effective technology with good clinical practicality. The use of HD-EEG offers several practical advantages to study brain function in NDDs such as ASD. Compared with MRI, EEG may have higher relative tolerance for movement, higher temporal resolution, and could be operated with a lower level of expertise at a considerably lower cost.

### Limitations

Our study has several limitations. First, methodological differences related to eye status (open/closed) must be considered. Local occipital overconnectivity in eyes-open states has indeed been found in several independent ASD cohorts[85, 98], and data from participants with eyes closed was recently shown to heavily impact regional homogeneity patterns, particularly in the occipital lobe[99]. Furthermore, a practical challenge in EEG source analysis is the widely used selection of a priori anatomic template to define the network nodes[100]. Nevertheless, our study is not safe from biases due to different parcellations of the cortex, and further work investigating the effects of template selection will be important to determine their generalizability[101]. Another problem when computing FC at the source level is the ‘spatial leakage’ as the reconstruction of true dipole sources from the scalp signals will be spread over numerous voxels. Here, the choice of the inverse solution/connectivity combination was supported by comparative studies using simulated data from a biophysical/physiological model[51], and real data recorded during a cognitive task[31]. Also, our study is blind to the physiological mechanisms underlying the present findings. One possibility might be that the developmental process of pruning of overabundant connectivity during early postnatal years is altered in ASD[102]. Deficient pruning of developing synaptic connections would lead to functional overconnectivity. More evidence will be required, however, to make strong claims about the physiological mechanism underlying local network overconnectivity in young children with ASD. Finally, although our results provide robust effect sizes on cortical network metrics, the sample size of our ASD group is relatively limited and we did not replicate our findings in an independent dataset. Further growth of public datasets in this particular age range has been challenging, but recent efforts will surely provide opportunity for cross-validation and stronger statistical power.

### Conclusions

In summary, here we used an innovative HD-EEG source reconstruction model to show that cortical network metrics in young children with ASD are characterized by specific FC overconnectivity patterns that contribute to distinctive ASD-related behavioral features. These results provide new insights into the complexity of altered brain function in ASD. Collectively, the stratification of well-defined neural signatures that give rise to the clinical heterogeneity of ASD has the potential to provide more accurate prognosis and help to select the optimal strategy for therapeutic intervention.

## Acknowledgements

The recruitment of participants for this research project was based on a solid synergy between the clinical and the research unit of our ASD center. We are really thankful to our clinical team for this close collaboration, which enables the identification of potential candidates and facilitates the recruitment and enrolment process. We also thank Laura Mendes and Katherina Gschwend for their help in the phenotypic assessment, as well as Romane Garcia for her expertise as a peer supporter concerned by ASD. We are deeply thankful to all children, their families and caregivers that are part of the STSA cohort for their participation in the study. We appreciate access to imaging data from the NIH Pediatric MRI database. This work was also possible thanks to the support of F. Hoffmann-La Roche.

## Availability of data and materials

The STSA dataset is available upon reasonable request to. The code used for the analysis is publicly available at https://github.com/stsa-research.

## Funding

This project was funded by F. Hoffmann-La Roche.

## Conflict of interest

The authors declare that they have no competing interests.

## Notes

### Competing Interest Statement

The authors have declared no competing interest.

### Summary of Updates

Add three authors due to their substantial contribution in the design and conception of the study.

