## Supplemental Material for "Social, cognitive and sensory dimensions of cortical network overconnectivity in young children with autism spectrum disorder"

Service des Troubles du Spectre de l'Autisme & apparentés

Centre Cantonal de l'Autisme

Les Allières, Al/05/508

Av. de Beaumont 23, CH-1011 Lausanne

The supplementary material has been provided by the authors to give readers additional information about their work.

**Supplementary methods**

1. **EEG PRE-PROCESSING WITH AUTOMAGIC TOOLBOX**

A channel was labelled as error-prone based on three criteria: i) when the correlation between the recorded data and an estimate of neighboring electrodes was r<0.85; ii) if it has more line noise relative to its signal than all other electrodes (4SD); and iii) if it exhibits a flatline for longer than 5 sec. Next, data were filtered using a 0.1Hz high-pass filter with a cutoff frequency of -6dB, and a ZapLine algorithm was applied to line noise artifacts^45^. To further remove noise contribution to the signal, electro-oculography (EOG) regression effectively mitigated artifacts caused by eye movements and separated them from the HD-EEG signal using the subtraction method that relies on linear transformation of the HD-EEG activity^46^, resulting in a reduction of the final number of channels to 118. After the artifact correction, residual bad channels were excluded if their SD exceeded a threshold of 20μV and/or if they showed a change of amplitude of ±80 μV at any temporal point. Subsequently, Automagic pipeline automatically assessed the quality of the resulting HD-EEG signals based on four criteria: i) if the rate of overall high-amplitude (OHA, >30 μV) data points in the signal was larger than 0.2; ii) if more than 20% of timepoints showed a variance larger than 15μV across channels; iii) if more than 30% of channels showed high variance (>15μV); and iv) if the rate of error-prone electrodes was higher than 0.15. In a next step, error-prone electrodes were removed and later interpolated using an EEGLab spherical spline interpolation.

2. **NETWORK METRICS**

Node-wise: The CC of a network can then be calculated as:

$$CC^{w}=\frac{1}{n}\sum_{i\in N} \frac{2t_{i}^{w}}{k_{i}^{w}\left( k_{i}^{w}-1 \right)}$$

Where $t_{i}^{w}$is the number of triangular connections around the node i:

$$t_{i}^{w}=\frac{1}{2}\sum_{j,h\in N} \left( w_{ij}w_{ih}w_{jh} \right)^{\frac{1}{3}}$$

BC is calculated as:

$$BC=\frac{1}{\left( n-1 \right)\left( n-2 \right)}\sum_{h,j\in Nh\neq j,h\neq i,i\neq j} \frac{\varphi_{hj}^{\left( i \right)}}{\varphi_{hj}}$$

Where $\varphi_{hj}$ is the number of shortest paths between node $h$ and node $j$, and $\varphi_{hj}^{\left( i \right)}$ is the number of shortest paths between $h$ and $j$ that pass-through node $i$.

LE is calculated as:

$$LE=\frac{1}{n\left( n-1 \right)}\sum_{j,k\in N} \frac{1}{L_{j,k}}$$

Where $L_{j,k}$ is the average distance (number of steps) between nodes $j$ and $k$ in the network.

Global topology: SWP is defined as:

$${SWP}^{w}=\frac{\frac{CC}{CC_{rand}}}{\frac{SPL}{{SPL}_{rand}}}$$

Where CC and $CC_{rand}$ are the clustering coefficients, and $SPL$ and ${SPL}_{rand}$ are the characteristic path lengths of the tested and the random network, respectively. EWCI is defined as the sum of the weights of the resultant subnetwork:

$$EWCI=\left( \sum_{e}^{M} w_{e} \right)\times100$$

where $w_{e}$ represents the weight of the edge $e$ in the subnetwork and $M$ is the number of edges.

3. **REGULARIZED CANONICAL CORRELATION ANALYSIS (RCCA)**

Five participants were excluded from the rCCA due to incomplete clinical data, leaving the final sample in 99 children (64 TD vs 35 ASD). The input to the rCCA were the 36 network-level dimensions from the list of ranked FC features and the following phenotypic measures: Verbal, nonverbal and full-scale IQ for the cognitive domain; the five SRS subscales T-scores (Awareness, Cognition, Communication, Motivation, Mannerisms) plus the total SRS T-score for the social domain; and the seven SPM subscales T-scores (Social, Visual, Hearing, Touch, Body, Balance, Planning) plus the total SPM T-score for sensory processing domain. The L2-penalty was settled for both variable sets (FC features and clinical measures) on a hyperparameter grid of λ_FC_ = [1,2,3,4,5] and λ_cog_ = [0.002,0.13], λ_srs_ = [0.002,0.02] and λ_sens_ = [0.002,0.2], respectively.

Before running the rCCA, we used the Spearman correlation between FC and the standard clinical measures in 50 subsampled replicates. We ranked FC features by how frequently they were selected as statistically significant, and used this rank list as input into rCCA to estimate sets of linear combinations of FC features that best relate to linear combinations of clinical features. We then generated a coarse grid of possible values for λ1 and λ2 and determine the optimal regularization parameters. Model performance was evaluated using a cross-validation approach, based on robust null permutation approach in 1,000 training set/test set replicates, with test data (15%) held out from feature selection and rCCA calculation.

**Supplementary figures**

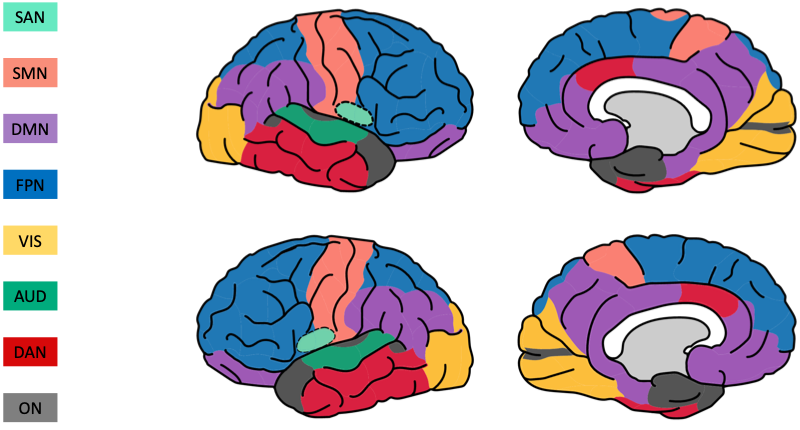

**Supplementary Figure S1.** Functional network parcellation of the brain. The cortical areas were parcellated with 68 seed regions, and these seed regions were assigned to 8 macroscale cortical networks: visual network (VIS), auditory network (AUD), sensorimotor network (SMN), fronto-parietal network (FPN), salience network (SAN), dorsal attentional network (DAN), default mode network (DMN) and other networks (OTHER). The corresponding information of each seed region is shown in Table S1.

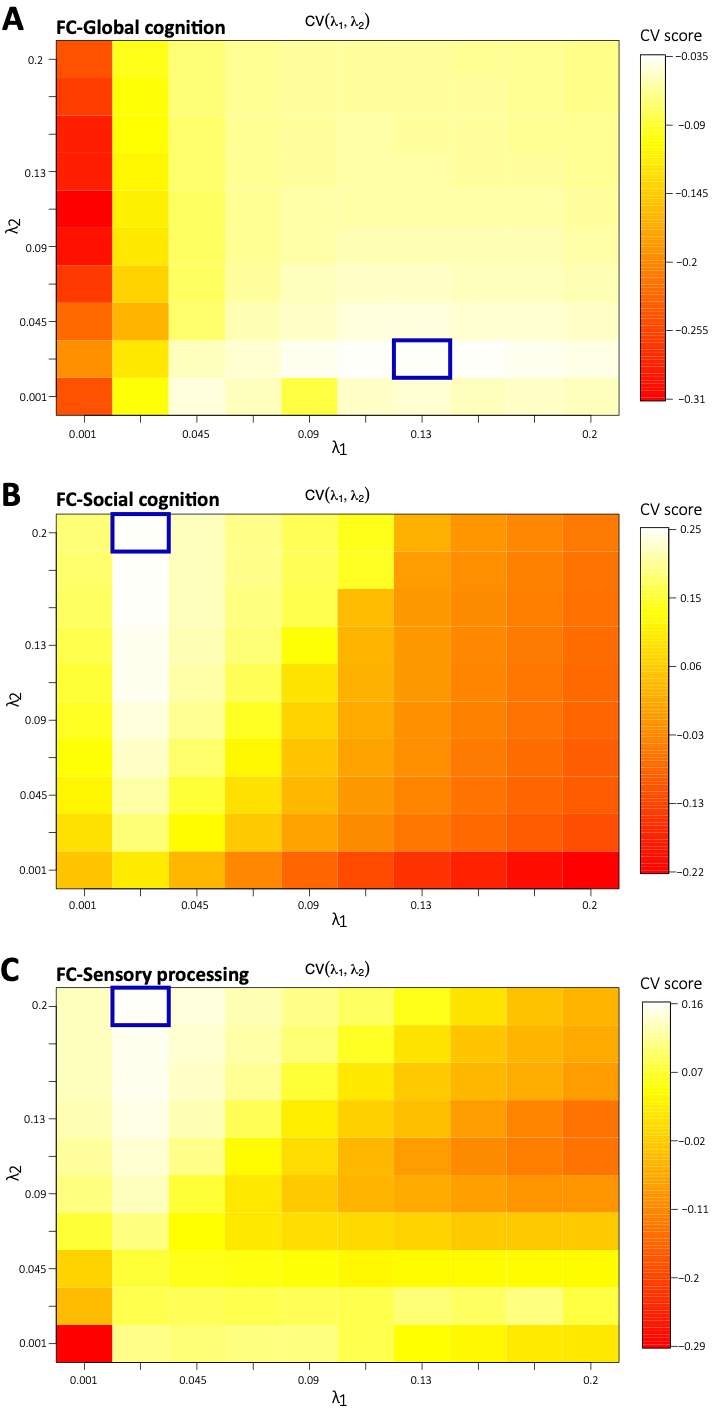

**Supplementary Figure S2.** Heatmaps used to select regularization parameters for the FC-Global cognition (**A**), FC-Social cognition (**B**) and FC-Sensory processing (**C**) rCCA. We used the Cross-Validation Ridge method, where the highest cross-validated score is desired, in a constrained range between 0.001 and 0.2 (length = 10) to yield the highest canonical correlation of the first variate.

**Supplementary tables**

**Table S1.** Affiliation of the 68 cortical regions –derived from the Desikan-Killiany atlas–, to the eight large-scale resting-state cortical networks

| **Network of Interest** | **Brain regions** |
| --- | --- |
| **Dorsal Attention Network (DAN)** | Caudalanteriorcingulate (L)  Inferiortemporal (L/R)  Middletemporal (L/R) |
| **Auditory Network (AUD)** | Superiortemporal (L/R) |
| **Visual Network (VIS)** | Cuneus (L/R)  Fusiform (L/R)  Lateraloccipital (L/R)  Lingual (L/R) |
| **Frontoparietal Network (FPN)** | Caudalmiddlefrontal (L/R)  Parsopercularis (L/R)  Parsorbitalis (L/R)  Parstriangularis (L/R)  Rostralmiddlefrontal (L/R)  Superiorfrontal (L/R)  Superiorparietal (L/R)  Frontalpole (L/R) |
| **Default Mode Network (DMN)** | Inferiorparietal (L/R)  Isthmuscingulate (L/R)  Lateralorbitofrontal (L/R)  Medialorbitofrontal (L/R)  Parahippocampal (L/R)  Posteriorcingulate (L/R)  Rostralanteriorcingulate (L/R)  Precuneus (L/R)  Supramarginal (L/R) |
| **Somatomotor Network (SMN)** | Paracentral (L/R)  Postcentral (L/R)  Precentral (L/R) |
| **Salience Network (SAN)** | Insula (L/R)  Caudalanteriorcingulate (R) |
| **Other** | Bankssts (L/R)  Entorhinal (L/R)  Pericalcarine (L/R)  Temporalpole (L/R)  Transversetemporal (L/R) |

**Table S2.** SRS-2 and SPM subscales T-scores of the study cohort of TD and ASD children expressed as Mean (SD).

|  |  | **TD (N=66)** | **ASD (N=38)** |
| --- | --- | --- | --- |
| **Social cognition (SRS-2)** | **Awareness** | 45.8 (7.8) | 64.3 (11) |
|  | **Cognition** | 45.2 (5.5) | 70.3 (9.1) |
|  | **Communication** | 45.6 (5.3) | 69.2 (10.4) |
|  | **Motivation** | 46.3 (6) | 64.7 (10.7) |
|  | **Mannerisms** | 45.1 (4.5) | 71.1 (12) |
| **Sensory processing (SPM)** | **Visual** | 50.6 (7.7) | 65.7 (8.1) |
|  | **Auditory** | 52.7 (8.7) | 63.3 (8.7) |
|  | **Tactile** | 51.9 (8.2) | 63.4 (10.7) |
|  | **Proprioception** | 51.7 (7.7) | 61.8 (8.5) |
|  | **Vestibular** | 49.1 (6.9) | 61.8 (9.1) |
|  | **Praxis** | 47.5 (6.8) | 65.4 (7.4) |
|  | **Social participation** | 49.8 (7.6) | 67.1 (7.8) |

ASD: Autism Spectrum Disorder, TD: Typically Developing, SRS-2: Social Responsiveness Scale-2, SPM: Sensory Processing Measure.

**p < 0.05; **p < 0.01; ***p < 0.001*
